# Atrial arrhythmogenesis in ex vivo aged mouse hearts with hypokalemia and right atrial stretch

**DOI:** 10.1101/2023.09.05.555978

**Authors:** Jessica Cayton, Zahra Nourian, Michelle Lambert, Zhenguo Liu, Timothy L. Domeier

## Abstract

**Introduction:** Atrial Fibrillation (AF) and atrial flutter (AFL) are the two most common cardiac arrhythmias in the United States. While advanced age has been correlated to AF/AFL, the lack of an appropriate animal model has hindered progress on better understanding the pathophysiology of atrial arrhythmogenesis. Both hypokalemic conditions and hemodynamic stretch have been associated with atrial tachyarrhythmias in patient populations. The purpose of this study was to examine the incidence of atrial tachyarrhythmias in an ex vivo aging C57BL/6 mouse model following hypokalemia and stretch challenges.

**Methods:** Hearts were isolated with combined cannulation of the aorta and superior vena cava in a modified right-sided working heart perfusion technique. Isolated hearts of Aged (26-29 month) male (n=14) and female (n=14) mice were subjected to normokalemic and hypokalemic conditions ± atrial preload elevation to 12 cmH_2_0 to induce atrial stretch. Heart rate, right ventricular (RV) pressure development, and incidence of atrial tachyarrhythmias were monitored using a pressure catheter and intracardiac electrocardiogram.

**Results:** In response to hypokalemia, there were no changes in mean heart rate, RV pressure development, or RV Rate-Pressure Product (Rate x RV peak pressure). Atrial tachyarrhythmias were not observed under baseline conditions, and only 1 of 8 hearts exhibited atrial tachycardia following the hypokalemia challenge. In response to atrial preload elevation, there was an increase in heart rate (P=0.0006 versus baseline) with no change in RV pressure development. RV Rate-Pressure Product was significantly elevated (P=0.013 versus baseline) with atrial preload due to the increase in heart rate.

Atrial tachyarrhythmias were not observed under both baseline conditions and following atrial preload elevation. In response to the combined hypokalemia and preload challenges, there was an increase in heart rate (P=0.008 versus baseline) with no change in RV pressure development or RV Rate Pressure product. Atrial tachyarrhythmias were not observed under baseline conditions, yet after the combined challenges 50% of aged hearts exhibited atrial tachycardia or AF/AFL. During bouts of AF/AFL, the AF/AFL led to a variable ventricular response and concomitant contractile dysfunction in the form of variable RV pressure development.

**Conclusion:** Ex vivo aged mouse hearts exhibit atrial tachyarrhythmias in response to combined hypokalemia and right atrial stretch conditions. The aged C57BL/6 mouse model is therefore useful for pre-clinical studies of atrial arrhythmogenesis.

## Introduction

Atrial tachyarrhythmias are the most prevalent cardiac arrhythmias with a significant and world-wide contribution to morbidity and mortality (Li et al., 2022; Orejarena et al., 1998; Vuong et al., 2019). The most common atrial arrhythmias are focal atrial tachycardias, atrial fibrillation (AF), and atrial flutter (AFL), and each can initiate with ectopic pacemaker formation(Chen et al., 1994; Lee et al., 2012; Saoudi et al., 2001). Focal atrial tachycardias often originate within a single site within the atrium (Kalman et al., 1998; Kistler et al., 2005; Kistler et al., 2003), and in atrial tissue with structural remodeling and/or fibrosis ectopic foci trigger macroreentrant reentry which manifests clinically as AF/AFL (Gong et al., 2007; Saoudi et al., 2001; Schotten et al., 2011; Stanley Nattel, 2003). While atrial tachycardia and AF are most seen in the left atria and AFL most common in the right, any of these tachyarrhythmias may occur in either atrium(Kistler et al., 2003; Li, 2022; Saoudi et al., 2001).

Atrial Fibrillation is defined by rapid *irregular* ectopic activity (Blume et al., 2011; De Jong et al., 2011; Moon et al., 2015; Rostock et al., 2008) and AFL characterized by rapid *regular* atrial activity (Waldo, 2001; Waldo & Feld, 2008), yet AFL can cardio-convert into AF making the two forms of arrhythmia difficult to distinguish (Granada et al., 2000; Li, 2022; Saoudi et al., 2001). Clinically, AF and AFL are similar in mortality rates(Waldo, 2001), and while AF/AFL can be present without symptoms the conditions commonly associate with cardiac palpitations, shortness of breath, and fatigue. In the long-term, AF/AFL leads to cardiac failure, stroke, and sudden death (Lip et al., 2016). As AF/AFL continues to increase in incidence and prevalence, these tachyarrhythmias are expected to become one of the largest epidemics and public health challenges within the next 30 years (Lippi et al., 2021).

The elderly are at greatest risk for AF/AFL with prevalence estimated at approximately 10% in individuals over the age of 80 (Go et al., 2001). The aged myocardium is subjected to many factors that can lead to arrhythmogenesis, and two of the strongest arrhythmia triggers include hemodynamic stretch and hypokalemic (low K^+^) conditions (De Jong et al., 2011; Krijthe et al., 2013; K. Tazmini et al., 2020). Despite the growing importance of AF/AFL in human health, biomedical research studies of spontaneous AF/AFL are limited, particularly in mouse models (Schuttler et al., 2020). Historically, the low incidence of AF/AFL in mouse models was attributed to the small size of the mouse atrium and limited tissue substrate to sustain AF/AFL (Byrd et al., 2005; Garrey, 1914; Janse & Rosen, 2006). While studies using novel mouse transgenic models have challenged this simplified view (Nishida et al., 2010; Riley et al., 2012; Schuttler et al., 2020), many such models exhibit severe atrial remodeling, aggressive pathological cardiac dysfunction, and decreased lifespan which may not translate to human pathophysiology, particularly in the early stages of disease progression. Furthermore, cardiac stretch is a well-established pro-ectopic factor in humans and large animals, yet remains difficult to study in mice in vivo due to high heart rates and limited atrial filling time. Therefore, to circumvent such challenges most investigations in mouse models use programmed electrical stimulation protocols to induce atrial arrhythmia (Schuttler et al., 2020), and this is not a typical ectopic trigger in humans.

This investigation examined incidence of right atrial arrhythmias in aged mouse hearts following ex vivo challenges, with the goal of establishing conditions to study spontaneous AF/AFL in the murine myocardium. We tested the hypothesis that hypokalemia and atrial stretch would lead to spontaneous tachyarrhythmias in isolated perfused mouse hearts of aged C57BL/6 mice.

## Methods

### Animal models

Animal procedures were approved by the Animal Care and Use Committee at the University of Missouri (Approval reference number 35701) and complied with all US regulations involving animal experiments. Mice utilized for this study included male and female C57BL/6 mice bred and raised at the University of Missouri and aged for 26-29 months.

### Cardiac Isolation and Experimental Procedure

Mice were placed under deep anesthesia via ketamine/xylazine intraperitoneal injection (100 mg/kg:5mg/kg). Once deep anesthesia was ensured via lack of pedal reflex, hearts were isolated and cannulated via the ascending aorta (within ∼3 minutes) and Langendorff perfused (60 mmHg, 37 ºC) with oxygenated (95% O_2_/5% CO_2_) Krebs– Henseleit buffer (KHB, *normokalemic*) which contained (in mM): 117 NaCl, 4.7 KCl, 1.2 MgSO_4_, 1.2KH_2_PO_4_, 25 NaHCO_3_, 11.11 Glucose, 0.4 Caprylic Acid, 1 Pyruvate, 0.5Na EDTA, and 1.8 CaCl_2_. The superior vena cava was cannulated for right-atrial preload perfusion, a Millar 1.0 F pressure catheter was placed into the right ventricle to obtain right ventricular pressure values, and an octapolar electrocardiogram (ECG) catheter was placed into the right atria to obtain atrial and ventricular electrical activity. Data were monitored and collected with LabChart software. After cannulation and stabilization of the preparation, hearts were perfused in Langendorff mode (0 cmH_2_0 applied preload) with normokalemic KHB for 10 minutes for baseline measurements. After baseline was obtained, hearts were subjected to an intervention of 1) hypokalemia challenge in langendorff mode (no stretch), 2) right atrial stretch challenge under normokalemic conditions, or 3) combined hypokalemia and right atrial stretch challenge. Hypokalemic conditions were obtained by changing all buffer in the system to 2 mM K^+^ by lowering KCl (to 1.0 mM) and KH_2_PO_4_ (to 1 mM). Right atrial stretch was induced by opening the preload cannula and perfusing the right atrium with oxygenated (95% O_2_/5% CO_2_) KHB at a constant pressure of 12 cmH_2_0. Following initiation of the respective challenges, pressure/ECG cannula were adjusted and hearts were allowed to re-equilibrate for 3-5 minutes prior to criterion measurements. This time was defined as post-intervention t=0.

### ECG analysis

In total, 8 hearts were studied with hypokalemia challenge, 8 hearts with atrial stretch challenge, and 15 hearts with the combined conditions. With combined conditions 3 of 15 hearts developed sustained ventricular fibrillation which precluded atrial tachyarrhythmia analysis, and therefore these hearts were excluded from final pressure and atrial arrhythmia analysis which decreased the total number analyzed to 12 in this group. Severe atrial tachyarrhythmias considered for this study included tachycardia and AF/AFL. Tachycardia was identified as high amplitude atrial ECG waveforms with rapid rate, and on average the tachycardic rate was 57.8 +/-0.22 percent higher than baseline sinus rhythm. AF/AFL was identified as a rapid atrial rate with variable ventricular response. Right ventricular rate and pressure values were analyzed for 2 minutes prior to intervention for baseline, and post-intervention t=0 to t=2. Tachyarrhythmia incidence was monitored during 10-minute baseline interval, during first post-intervention interval of t=0 to t=10, and second post-intervention interval of t=10 to t=20. R-R intervals (elapsed time between two successive ventricular ECG deflections) were measured during bouts of AF/AFL that lasted longer than one second and compared with one second intervals in the preceding sinus rhythm.

### Data presentation and statistics

Right ventricular rate and pressure data were analyzed using paired t tests (baseline versus challenge). Arrhythmia data were analyzed using Fisher’s Exact test at 0-10 and 10-20 minutes after intervention, compared to 10 minutes of baseline. F-test was used to calculate differences in measurement variance. A p value of less than 0.05 was considered statistically significant.

## Results

The goal of this investigation was to examine factors contributing to atrial tachyarrhythmias in the aged murine myocardium. The first factor examined was hypokalemia (2 mM K^+^, ***Figure 1A-B***). Following baseline measurements in langendorff perfused heart mode at 6 mM external K^+^, K^+^ was lowered to 2 mM for 20 minutes while heart rate (***Figure 1C***), RV pressure (***Figure 1D***), RV Rate-Pressure product (***Figure 1E***), and incidence of atrial tachycardia/AF/AFL was monitored (***Figure 1F***). With hypokalemia, there were no changes in mean heart rate (***Figure 1C***), RV pressure development (***Figure 1D***), or RV Rate-Pressure product (***Figure 1E***). Atrial tachyarrhythmias were not observed under baseline conditions, and only 1 of 8 hearts exhibited atrial tachycardia following the hypokalemia challenge (***Figure 1F***).

**Figure 1:**
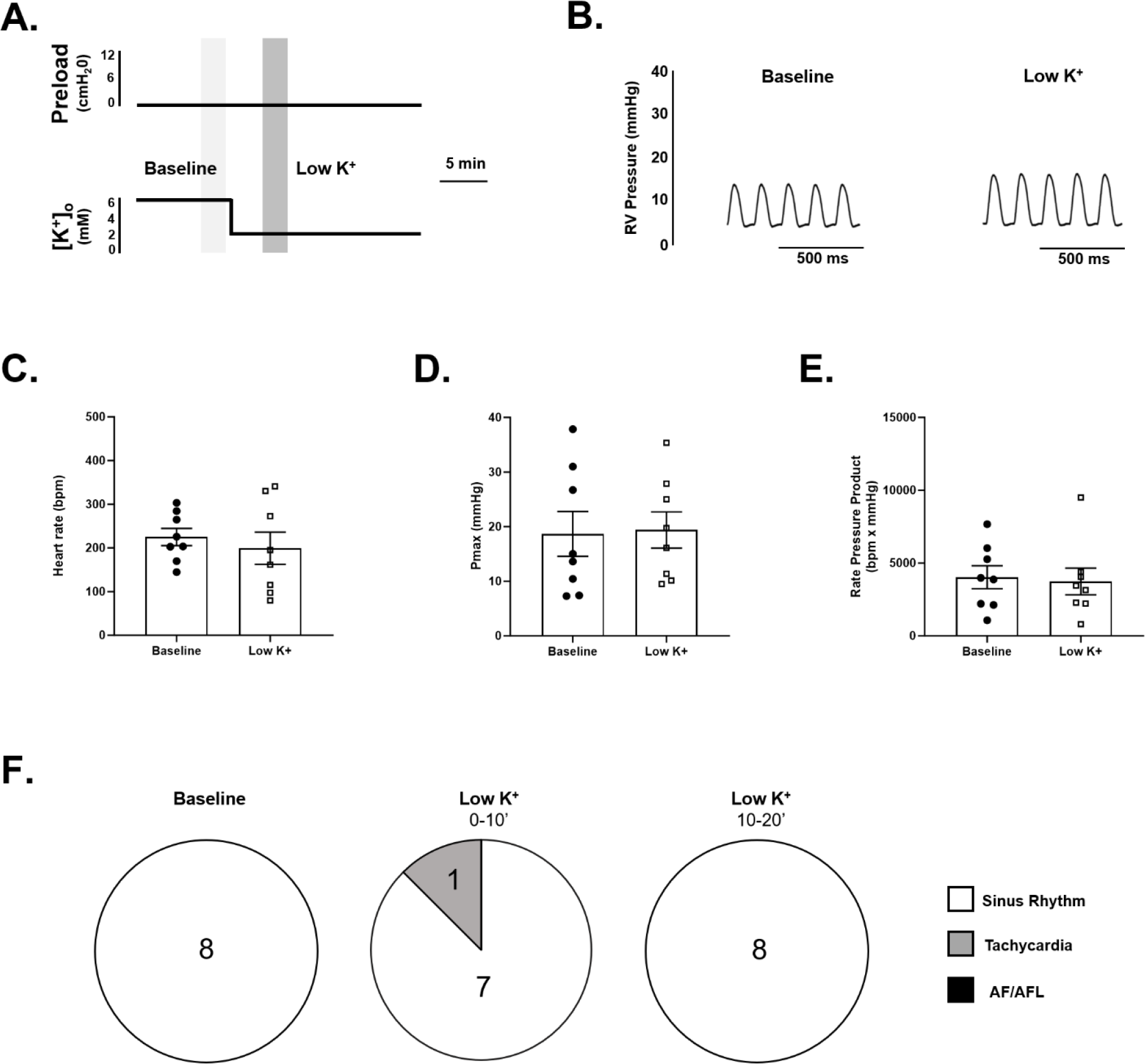
Effect of hypokalemia on heart rate, right-ventricular function, and atrial tachyarrhythmias in aged hearts. **A**) Protocol for hypokalemia (2 mM extracellular K^+^) challenge of isolated hearts of aged mice. **B)** Example right-ventricular pressure development under baseline langendorff conditions (left panel) and following hypokalemia challenge (low K^+^, right panel). **C-E)** Summary data of heart rate (**C**), maximum pressure development (Pmax, **D**), and rate-pressure product (RPP, **E**) under baseline langendorff conditions (closed circles) and following hypokalemia challenge (open squares). **F)** Summary data of most severe atrial arrhythmia observed under 10 minutes of baseline conditions (left), 0-10 minutes following hypokalemia challenge (center), and 10-20 minutes following hypokalemia challenge (right). Number of hearts with each condition (sinus rhythm or atrial tachycardia) listed within pie chart. P=0.40 versus baseline (**C**); P=0.57 versus baseline (**D**); P=0.72 versus baseline (**E**). n=8 (3 male/5 female, 27-29 months of age), paired t-test.

The second pro-arrhythmogenic factor examined was right atrial preload elevation (***Figure 2A-B***) and atrial stretch. Following baseline measurements in Langendorff perfused heart mode (i.e., 0 cmH_2_O with no applied atrial preload), atrial preload was elevated to 12 cmH_2_O for 20 minutes while heart rate (***Figure 2C***), RV pressure (***Figure 2D***), RV Rate-Pressure product (***Figure 2E***), and incidence of atrial tachycardia/AF/AFL was monitored (***Figure 2F***). In response to atrial preload elevation, there was a significant increase in heart rate (***Figure 2C***) with no change in RV pressure development (***Figure 2D***). RV Rate-Pressure product was also significantly elevated (***Figure 2E***) due to the increase in heart rate. Atrial tachyarrhythmias were not observed under both baseline conditions and following atrial preload elevation (***Figure 2F***).

**Figure 2:**
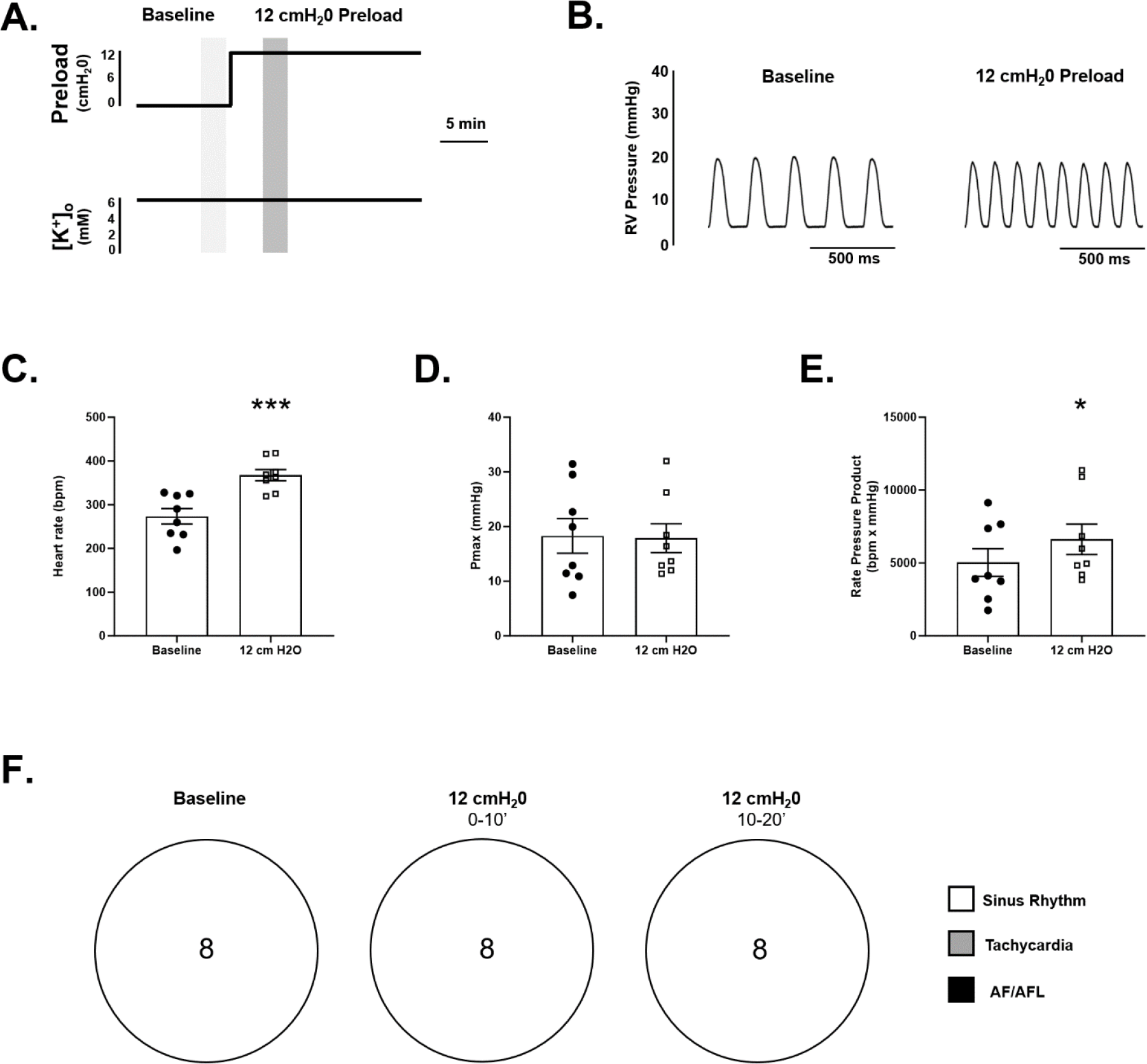
Effect of right-atrial preload increase on heart rate, right-ventricular function, and atrial tachyarrhythmias in aged hearts. **A**) Protocol for right atrial preload challenge (12 cmH_2_O) of isolated hearts of aged mice. **B)** Example right-ventricular pressure development under baseline Langendorff conditions (left panel) and following right-atrial preload elevation to 12 cmH_2_O (right panel). **C-E)** Summary data of heart rate (**C**), maximum pressure development (Pmax, **D**), and rate-pressure product (RPP, **E**) under baseline langendorff conditions (closed circles) and following preload elevation to 12 cmH_2_0 (open squares). **F)** Summary data of most severe atrial arrhythmia observed under 10 minutes of baseline conditions (left), 0-10 minutes following 12 cmH_2_O right atrial stretch (center), and 10-20 minutes following 12 cmH_2_O right atrial stretch (right). *** P=0.0006 versus baseline (**C**); P=0.84 versus baseline (**D**); * P=0.013 versus baseline (**E**), n=8 (4 male/4 female, 26-29 months of age), paired t-test.

The final set of experiments examined the combined challenges of hypokalemia with atrial preload. Following baseline measurements in Langendorff perfused heart mode (i.e., 0 cmH_2_O and 6 mM K^+^), atrial preload was elevated to 12 cmH_2_O with hypokalemia for 20 minutes (***Figure 3A-B)***. In response to the combined challenge, there was a significant increase in heart rate (***Figure 3C***) with no change in RV pressure development (***Figure 3D***) or RV Rate-Pressure product (***Figure 3E***). Atrial tachyarrhythmias were not observed under baseline conditions, yet after the combined challenge 50% of aged hearts exhibited atrial tachycardia or AF/AFL (***Figure 3F***). Observed atrial arrhythmias were present in hearts of both male and female mice (4 male, 3 female). During bouts of AF/AFL, the AF/AFL led to a variable ventricular response and concomitant cardiac dysfunction in the form of variable RV pressure development (***Figure 4***). Taken together, these data indicate that combined hypokalemia and atrial preload challenge induce atrial tachyarrhythmias in isolated hearts of aged mice.

**Figure 3:**
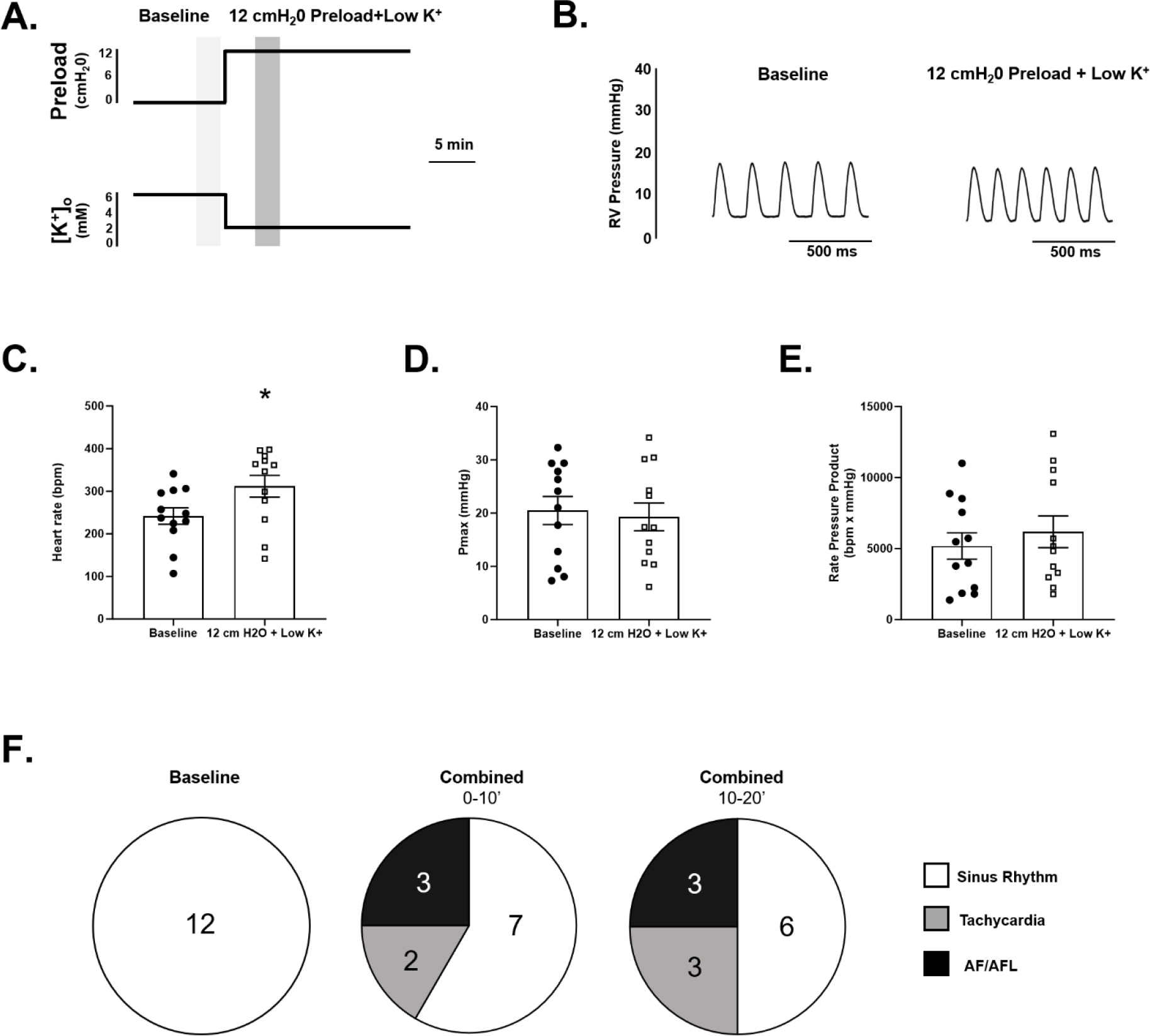
Effect of combined hypokalemia and right-atrial preload challenge on heart rate, right-ventricular function, and atrial tachyarrhythmias in aged hearts. **A**) Protocol for hypokalemia challenge (2 mM extracellular K^+^) and right atrial preload challenge (12 cmH_2_O) of isolated hearts of aged mice. **B)** Example right-ventricular pressure development under baseline langendorff conditions (left panel) and following combined hypokalemia challenge and right-atrial preload elevation to 12 cmH_2_O (right panel). **C-E)** Summary data of heart rate (**C**), maximum pressure development (Pmax, **D**), and rate-pressure product (RPP, **E**) under baseline langendorff conditions (closed circles) and following combined hypokalemia challenge and preload elevation (open squares). **F)** Summary data of most severe atrial arrhythmia observed under 10 minutes of baseline conditions (left), 0-10 minutes following combined hypokalemia challenge and 12 cmH_2_O right atrial stretch (center), and 10-20 minutes following combined hypokalemia challenge and 12 cmH_2_O right atrial stretch (right). Number of hearts with each condition (sinus rhythm, atrial tachycardia, AF/AFL) listed within pie chart. Atrial tachycardia/AF/AFL incidence with combined hypokalemia challenge and 12 cmH_2_O right atrial stretch was greater than paired baseline during both 0-10 minute interval (**P=0.0186) and 10-20 minute interval (***P=0.0069). One-tailed Fisher’s exact test. Atrial tachycardias were observed in both males and females (n=2 males/1 female). A total of 4 hearts exhibited AF/AFL during the protocol (n=2 males/n=2 females), with one heart cardioverting from AF/AFL (0-10’) to Tachycardia (10-20’). * P=0.013 versus baseline (**C**), P=0.60 versus baseline (**D**); P=0.27 versus baseline (**E**), n=12 (7 male/5 female, 26-29 months of age), paired t-test.

**Figure 4:**
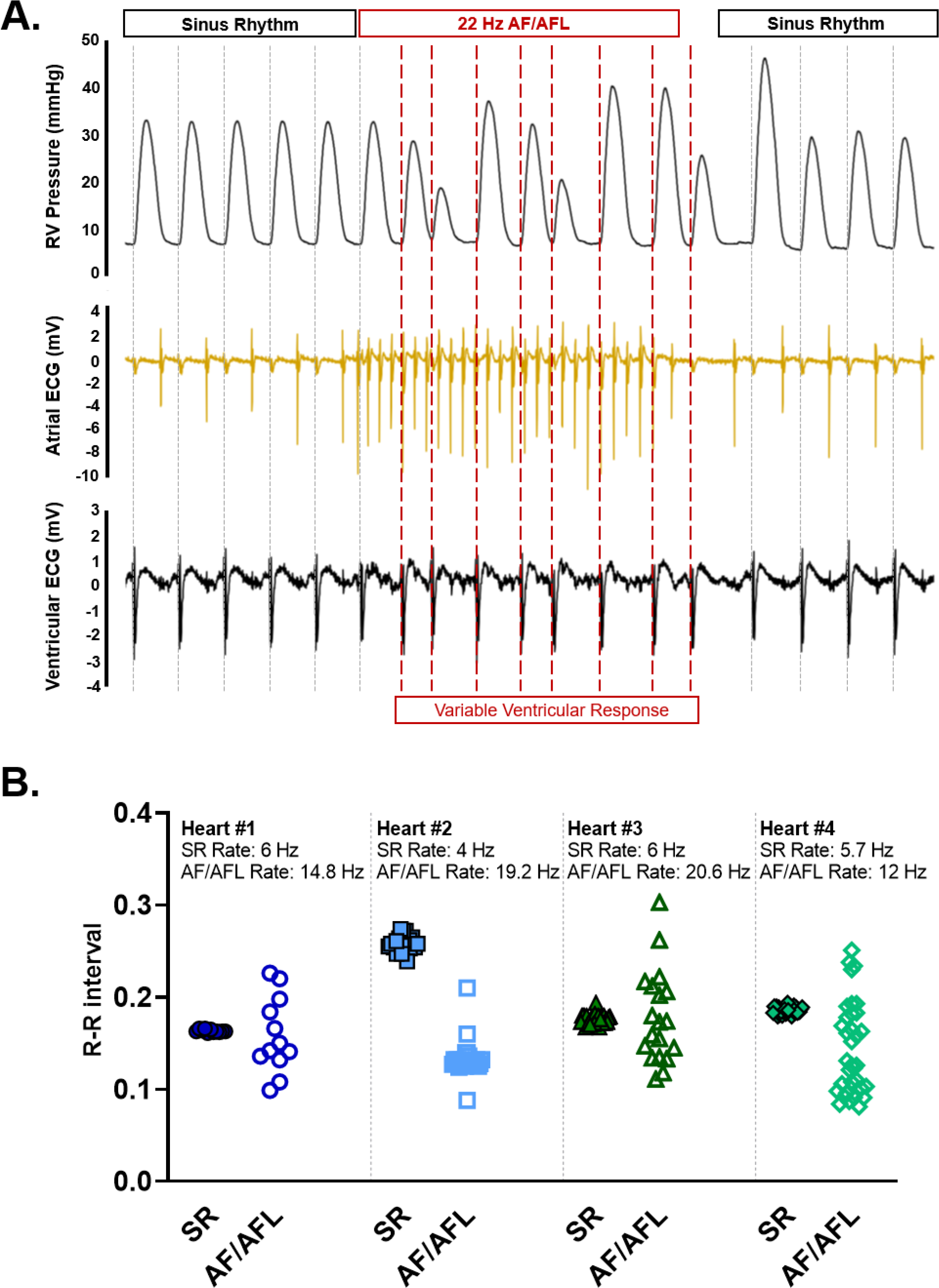
Variable ventricular response during atrial fibrillation/flutter in aged hearts. **A)** Example traces of right ventricular pressure (upper panel), Atrial ECG (middle panel), and Ventricular ECG (lower panel) during sinus rhythm (SR) and 22 Hz atrial fibrillation/flutter (AF/AFL) during combined hypokalemia and atrial stretch conditions. Dashed lines indicate ventricular response. **B**) Comparison of R-R interval during SR and AF/AFL in four hearts during combined hypokalemia and atrial stretch conditions. Average SR rate and AF/AFL rate for each heart listed above raw data points. Within each heart, R-R interval had equal variance during SR (*F* test P>0.05) and unequal variance during AF/AFL (*F* test P=0.0001 Hearts 1, 3, and 4; *F* test P=0.0002 Heart 2).

## Discussion

Atrial tachyarrhythmias are a growing health problem worldwide, yet the mechanisms underlying arrhythmogenesis are still not fully understood. The objective of this study was to examine the effect of two known risk factors, hypokalemia and atrial stretch, on atrial arrhythmias in ex vivo mouse hearts. Our main finding was that the combination of both conditions was necessary to induce spontaneous atrial tachyarrhythmias in aged C57BL/6 mouse hearts. As discussed in recent comprehensive reviews, murine models offer unique opportunities to investigate atrial tachyarrhythmias including AF (Keefe et al., 2022; Riley et al., 2012; Schuttler et al., 2020). However, very few mouse models exhibit spontaneous tachyarrhythmias and the majority require supraphysiological stimuli to induce arrhythmogenesis (Keefe et al., 2022). Atrial tachyarrhythmias are often intermittent in nature which hinders detection in vivo, particularly without concomitant triggering factors. In the absence of spontaneous arrhythmias in murine models, burst pacing protocols are experimentally used to trigger arrhythmias and examine the structural substrate (e.g., hypertrophy and fibrosis) present which sustains episodes of AF/AFL (Riley et al., 2012). While such protocols clearly probe AF/AFL substrate, they may be less ideal to examine potential ectopic triggers of AF/AFL (Riley et al., 2012).

A major advantage of the ex vivo approach in this study is induction of arrhythmias with experimental challenges associated with human risk factors for atrial tachyarrhythmias. Hypokalemia has proven to be a strong driver of arrhythmogenesis in the ventricle (Osadchii, 2010), in atrial and ventricular cardiomyocytes (K. Tazmini et al., 2020), as well as in human AF patients (Krijthe et al., 2013). In ventricular myocytes and atrial myocytes with organized t-tubule structure, reduced extracellular potassium levels promote calcium overload and associated electrical disturbances including early and delayed afterdepolarizations (Kiarash Tazmini et al., 2020). Interestingly, in atrial myocytes that lack t-tubules, hypokalemia may be arrhythmogenic via distinct mechanisms such as sodium current reactivation and early-afterdepolarizations (Kiarash Tazmini et al., 2020). Dietary potassium restriction and resulting hypokalemic conditions can be obtained in mice in vivo (Grune et al., 2022), yet without the precise experimental control of potentially additive risk factors including stretch.

Left-atrial volume index in humans is an independent predictor of AF (Leung et al., 2010), and atrial myocyte stretch is known to trigger arrhythmias (Kalifa et al., 2003; Leung et al., 2010; Nazir & Lab, 1996; Solti et al., 1989). One proposed mechanism of stretch-induced atrial arrhythmias is activation of stretch-activated non-selective cation channels (Bode et al., 2000) which promotes AF through calcium overload, spontaneous calcium release via RyR channels, and delayed afterdepolarizations (Dobrev & Wehrens, 2017; Nattel et al., 2020). Stretch-induced ectopic activity is difficult to trigger in vivo in rodent models due to the rapid heart rate and limited chamber filling time (Nishida et al., 2010; Schuttler et al., 2020). In the ex vivo preparation, baseline heart rate was 200-300 beats per minute which is approximately half of what is expected in vivo (Ho et al., 2011). From this baseline, atrial stretch caused an increase in heart rate (*Figure 2*) consistent with the classic Bainbridge effect (Hakumäki, 1987). Interestingly, we did not observe atrial tachyarrhythmias under stretch conditions alone, which may be in part to the increased heart rate with stretch and resulting overdrive suppression of ectopic pacemaker activity. However, combined atrial stretch and hypokalemia challenge induced spontaneous atrial tachyarrhythmias, and these arrhythmic bouts were intermittent and of variable duration consistent with prior studies in aged mice (Jansen et al., 2017). Our study is also in agreement with previous studies in animal models where combined risk factors associate with arrhythmogenesis (Chelu et al., 2009; Grune et al., 2022; McCauley et al., 2020).

This study specifically examined hearts of aged mice due to the high prevalence of arrhythmias in the elderly and the unique structure and function of the aged versus young myocardium. Established changes to the aged myocardium include increased mitochondrial oxidative stress, dysregulated autophagy, increased fibrosis, altered molecular signaling pathways and ion channel function, aberrant electrical pathways, and atrial hypertrophy and chamber dilation (Guo et al., 2014; Hamilton & Terentyev, 2019; Jansen et al., 2017; Obas & Vasan, 2018). In addition, impaired sinus node function due to a reduction in the number of sinus node pacemaker myocytes (Moslehi et al., 2012), slower firing rate of sinoatrial myocytes (Larson et al., 2013; Sharpe et al., 2017), and increased fibroblasts and elastin within atrial tissue (Moslehi et al., 2012; Sharpe et al., 2017) reduces automaticity and extends the interval during which latent pacemakers can drive cardiac excitation. Together, such changes predispose the aged atrium to both ectopic arrhythmia triggers, and provide the tissue-level structural substrate necessary to sustain complex tachyarrhythmias.

## Conclusions

The main finding of the present study was that a combination of hypokalemia and right atrial stretch in ex vivo aged C57BL/6 mouse hearts triggered spontaneous atrial tachyarrhythmias. This methodology will serve as an ideal experimental platform to test multiple factors related to atrial tachycardia and AF/AFL in future investigations. Mouse models are therefore useful for pre-clinical testing of pharmacological or gene therapy approaches to prevent ectopic triggers for atrial arrhythmias including AF/AFL when utilizing ex vivo techniques.

## Notes

### Competing Interest Statement

The authors have declared no competing interest.

